# Dual Role of Plasmacytoid Dendritic Cells in Humoral and CD8⁺ T Cell Memory Post COVID-19 mRNA Vaccination

**DOI:** 10.1101/2025.06.13.659539

**Authors:** Chiara Pizzichetti, Irene Latino, Tommaso Virgilio, Arianna Capucetti, Kamil Chahine, Luciano Cascione, Simone G. Moro, Melanie Brügger, Nedim Kozarac, Alain Pulfer, Louis Renner, Roshan S. Thakur, Charaf Benarafa, Daniel F. Legler, Santiago F. González

## Abstract

The Pfizer-BioNTech coronavirus vaccine (BNT162b2), one of the first nanoparticle-based vaccines approved by the World Health Organisation (WHO), demonstrated 95% efficacy in preventing against Severe acute respiratory syndrome coronavirus 2 (SARS-CoV-2) infection. However, the precise mechanism of action underlying its effectiveness remains poorly understood. This study investigated the early immune responses in the draining lymph node (dLN) and its role in mediating antiviral protection following vaccination. Here, we focused on the involvement of antigen-presenting cells (APCs) in adaptive immunity. In this study, we demonstrated that the Pfizer-BioNTech coronavirus vaccine is rapidly transported to the dLN and is primarily captured by leukocytes that initiate the expression of the viral antigenic spike protein. Notably, we demonstrated that plasmacytoid dendritic cells (pDCs) are key orchestrators of the inflammatory and humoral response, as their specific depletion led to impaired antibody production and diminished neutralization capacity.

Furthermore, single-cell transcriptomic analysis revealed an interaction between pDCs and CD8^+^ T cells that facilitates T cell activation. *In vivo* experiments confirmed that pDCs expressing the viral spike protein directly engage with CD8^+^ T cells, promoting their differentiation and expansion. Moreover, the absence of pDCs affected the formation of antigen-specific memory T cells. Overall, these findings highlight that pDCs are essential players in mediating both adaptive and humoral responses to the Pfizer-BioNTech coronavirus vaccine, providing insights into the mechanistic functioning of mRNA vaccines and establishing a novel role for pDCs as professional APCs.

## Introduction

Severe acute respiratory syndrome coronavirus 2 (SARS-CoV-2) was first detected in Wuhan, China, in December 2019^1^. According to the World Health Organization (WHO), it is the most recent large-scale pandemic, resulting in over 7 million deaths worldwide^2,3^. Various therapeutic approaches have been tested to prevent disease progression. However, despite extensive public health measures, the high transmissibility and mutational rate of the virus have continued to pose challenges in controlling the pandemic^4–6^.

While most patients were asymptomatic or showed mild symptoms, around 15% progressed to severe pneumonia, and 5% developed acute respiratory distress symptoms or septic shock^7,8^. For this reason, the WHO drew up guidelines to expedite and optimize the treatment of the disease. Antiviral medications like Remdesivir or Nirmatrelvir/Ritonavir and for a time monoclonal antibodies were recommended as early therapy to control virus replication, while corticosteroids, IL-6 receptor blockers, or Janus kinase inhibitors were used to mitigate excessive inflammatory responses in hospitalized patients requiring respiratory support^8,9^.

The emergence of symptomatic coronavirus disease 2019 (COVID-19) across the planet triggered a global race for the development of an effective vaccine. Building on existing discovery and preclinical data from SARS-CoV and MERS-CoV vaccine studies, close to 200 vaccine candidates were developed within months including recombinant protein-based vaccines, inactivated and live-attenuated SARS-CoV-2-based vaccines, various vector-based platforms, and innovative nucleic acid-based approaches^10,11^. Among these, RNA vaccines showed remarkable results in preclinical studies and accelerated clinical trials, and two of them, the Comirnaty-BNT162b2 (Pfizer-BioNTech) and the Spikevax (Moderna), were approved and have been used extensively^10,12–14^. BNT162b2 was the first RNA vaccine authorized, demonstrating 95% efficacy in preventing COVID-19^15,16^. Since then, Pfizer-BioNTech has released updated RNA vaccines to address the need for protection against new circulating SARS-CoV-2 variants^17^. However, despite its success, the immunological mechanisms driving the efficacy of these newly approved therapies remain incompletely elucidated^18^.

The principal route of administration for most vaccines, including the Pfizer-BioNTech coronavirus vaccine, is intramuscular injection. Following administration, the vaccine can reach the draining lymph node via lymphatic drainage or intracellular transport associated with antigen-presenting cells^19^. The specific pathway of lymph node access is known to shape the immune response, influencing both the magnitude and quality of immune cell activation and targeting ^20,21^. While clinical data have demonstrated that Pfizer-BioNTech administration induces local inflammation, lymphocyte infiltration, and spike protein expression at the injection site ^22–24^, the precise mechanism driving vaccine trafficking to the lymph node remains unclear. Moreover, recent studies have analyzed the immune response to the BNT162b2 vaccine in humans and mice. Kariko and colleagues discovered that the m1Ψ-modification on RNA reduces the sensing of the mRNA by most Toll-like receptors^25,26^. However, the vaccine successfully activated the melanoma differentiation-associated protein 5 (MDA5) in macrophages and dendritic cells. This resulted in triggering the interferon type I receptor (IFNAR1) signaling pathway, which is needed to activate the adaptive immune system^27^.

This study aims to dissect the early immunological events following Pfizer-BioNTech coronavirus vaccination, focusing on the role of pDCs in orchestrating both humoral and cell-mediated immune responses. We demonstrated that pDCs are the most efficient antigen-presenting cells (APC) expressing the spike in the draining lymph node (dLN). Additionally, the inflammatory cytokines released by pDCs contribute to the activation of other APCs and support the production of virus-specific IgG. Importantly, we showed that pDCs directly interact with CD8^+^ T cells, promoting their activation and memory differentiation. Our findings highlight pDCs as central players in linking innate to adaptive immunity and provide mechanistic insights in the mechanism of action of mRNA-based vaccines.

## Results

### Pfizer-BioNTech coronavirus vaccine is rapidly transported to the draining lymph node, accumulates in the medullary region, and initiates an IFN-mediated inflammatory response

To assess both the local and the systemic immune responses against the BNT162b2 vaccine, we immunized C57BL/6 mice via subcutaneous (s.c.) injection in the footpad (**Fig. 1A, Suppl. Fig. 1A**). We administered a dosage of 2μg/mouse, which has been shown to elicit immune responses comparable to those observed in humans^28^. The transport dynamics of the vaccine in real-time from the injection site to the draining popliteal lymph node (pLN) were analyzed by two-photon intravital microscopy. We observed a rapid arrival (of the labeled vaccine 10 min post-vaccination (p.v.)), which peaked at 30 min p.v. (**Fig. 1C)**, and accumulated primarily in the interfollicular (IF) and medullary regions of the pLN (**Fig. 1B, D; Suppl. Movie 1**). Subsequently, we conducted a time course study, monitoring the vaccine capture and the expression of the spike protein in leukocytic and non-leukocytic cells by flow cytometry. Remarkably, vaccine uptake and spike protein expression were already detectable 3 h p.v., with vaccine capture peaking at 12 h p.v. while the percentage of spike-expressing cells continued to increase up to 48 h p.v. respectively (**Fig. 1E, F; Suppl. Fig. 1B, C**). The analysis of CD45^−^ and CD45^+^ cells showed similar dynamics, with leukocytes being the primary contributors to vaccine uptake and spike protein expression (**Fig. 1E, F**). Next, we investigated the innate immune response by monitoring inflammatory and antibody production following vaccination. In line with previous studies^28^, our model induced a strong inflammatory response, correlating with an increase in the early expression of interferon signaling at 24 h p.v. (**Fig. 1G**). Furthermore, the vaccine triggered a robust local and systemic humoral response, marked by a significant increase in the anti-SARS-CoV-2 spike receptor binding domain (RBD) IgM titers at 7 and 10 days p.v **(Suppl. Fig. 1D, E**) and the IgG levels at 10 days p.v. (**Fig. 1H, I**) compared to non-vaccinated controls.

**Fig. 1.**
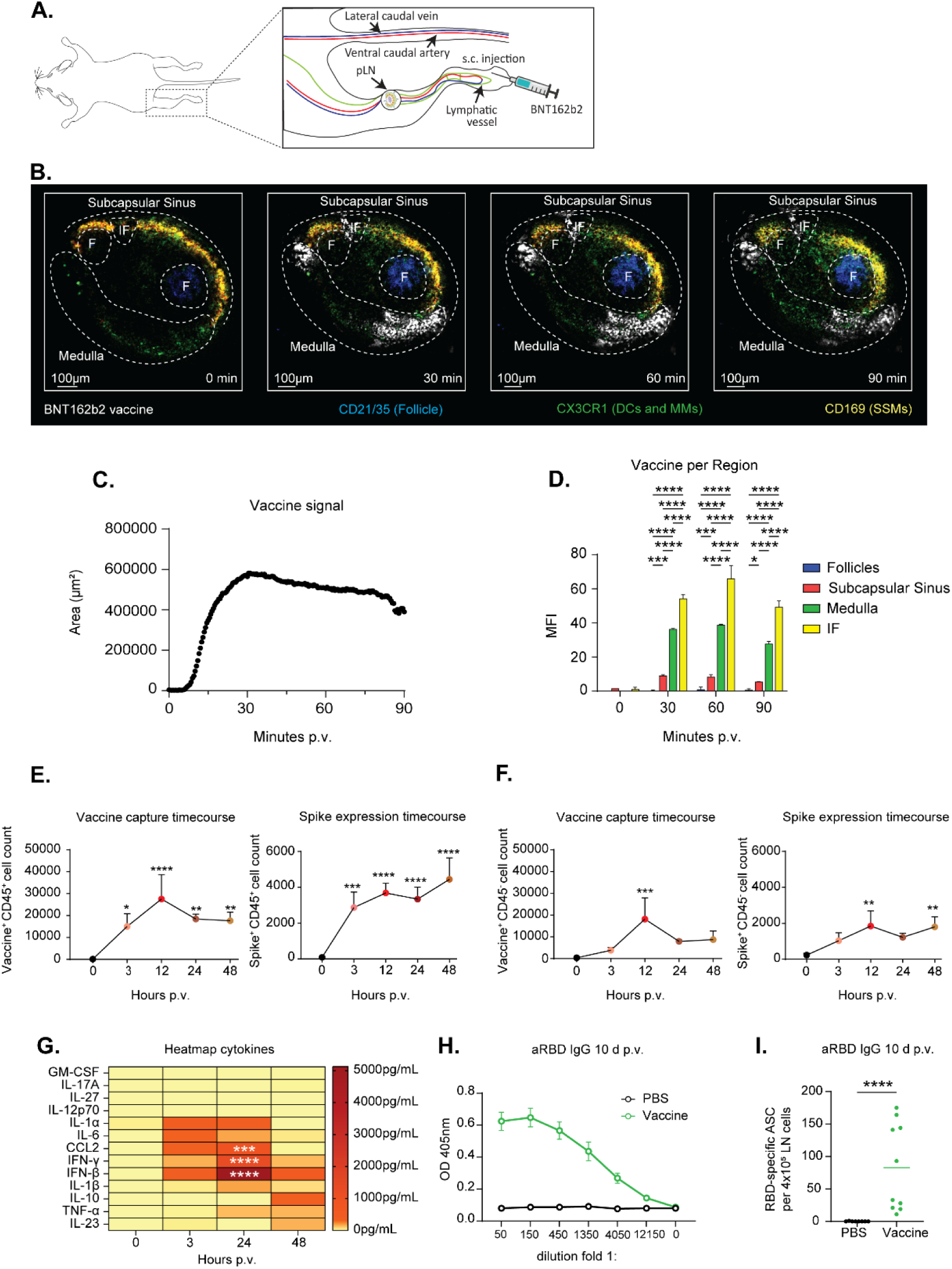
BNT162b2 vaccine has a rapid arrival in the dLN, where it gets captured by immune cells. **(A)** Experimental model of subcutaneous (s.c.) injection of the BNT162b2 vaccine in the footpad. **(B)** Representative intravital 2-photon micrographs acquired at 0, 30, 60, and 90 min p.v. IF stands for Interfollicular region; DCs, Dendritic cells; MMs, Medullary Macrophages; SSM, Subcapsular Sinus Macrophages. **(C)** Time course showing the arrival of the BNT162b2 vaccine in the pLN during the first 90 min p.v. **(D)** Quantification of the mean fluorescence intensity (MFI) of the vaccine in the different areas of the LN at 0, 30, 60, and 90 min p.v. **(E)** Capture of labeled BNT162b2 vaccine (left) and expression of coronavirus spike protein (right) by immune cells in the pLN at indicated time points p.v. **(F)** Capture of labeled BNT162b2 vaccine (left) and expression of coronavirus spike protein (right) by not immune cells in the pLN at indicated time points p.v. **(G)** Heatmap showing the lymph expression of different cytokines in the pLN at different time points p.v. **(H)** Anti-SARS-CoV-2 RBD IgG titers measured by ELISA at day 10 p.v. **(I)** Local anti-SARS-CoV-2 RBD IgG response measured by ELISPOT at day 10 p.v. **(E, F)** n=3 for 0 group, n=6 for all the other groups. Data are presented as mean ± SD. Two-way ANOVA **(D, G)** and One-way ANOVA **(E, F)** were followed by Bonferroni correction for multiple comparisons **(I)** Mann-Whitney U test. (* p < 0.05, ** p < 0.01, *** p < 0.001 **** p<0.0001).

### Plasmacytoid Dendritic Cells contribute to the humoral response to BNT162b2 vaccination

To further characterize the immune cell types involved in vaccine uptake and antigen expression, we performed a time course analysis in the pLN. Our study revealed that, among the cells capturing the vaccine, T and B cells were the most abundant (**Suppl. Fig. 2A, left panel**). Instead, analysing the cells expressing the spike protein, the APC, particularly dendritic cells (DCs), represented the predominant cell population (**Fig. 2A, right panel**). To determine the specific role of the different phagocytic populations, we analyzed the antibody response in vaccinated animals depleted of conventional DCs (cDCs) (*CD11c-DTR*) or macrophages (*CD169-DTR*). Given the critical role of the interferon (IFN) pathway in triggering an efficient humoral response to vaccines^29^, we also included a group of mice lacking the IFN type I receptor (IFNAR). We observed that DC depletion, but not macrophage depletion, was associated with a significant reduction in the systemic (**Fig. 2B**) and the local (**Fig. 2C**) antibody production, like that observed in *IFNAR knockout (KO)* mice. To further explore the specific contribution of different DC subsets to vaccine-induced immunity, we studied vaccine capture at 12 h p.v. and spike expression at 48 h p.v. in the three major groups of DCs (B220^+^ PDCA^+^ pDC, CD11b^+^ cDCs and CD11b^−^ cDCs). Notably, pDCs exhibited the highest percentage of cells positive for the vaccine and the spike protein at the evaluated time points (**Fig. 2D**). To confirm these results using a spatial imaging approach, we performed confocal microscopy analysis of sections at 12 h p.v., observing a pronounced accumulation of vaccine^+^ spike^+^ pDCs, in the medulla and the T cell zone of the draining LN (**Fig. 2E, F**). To further define the role of pDCs in the response to the vaccine, we assessed the levels of interferon response in mice depleted of pDCs (*aPDCA*), cDCs (*CD11c-DTR)*, macrophages (*CD169-DTR)*, and in *IFNAR KO* mice at 12 h p.v.. Interestingly, pDC-depleted mice, but not cDC- or macrophage-depleted mice, showed a reduction in IFN-β and IFN-γ protein levels, comparable to those observed in *IFNAR KO* mice (**Fig. 2G**). Furthermore, mice depleted of pDCs exhibited a significant reduction in the anti-SARS-CoV-2 RBD IgG levels following vaccination (**Fig. 2H**). In addition, we observed a reduced production of anti-SARS-CoV-2 RBD IgM on day 7 (**Suppl. Fig. 2B**), as well as a lower generation of anti-SARS-CoV-2 RBD IgG and RBD-specific memory B cells on day 10 p.v. (**Fig. 2I**).

**Fig. 2.**
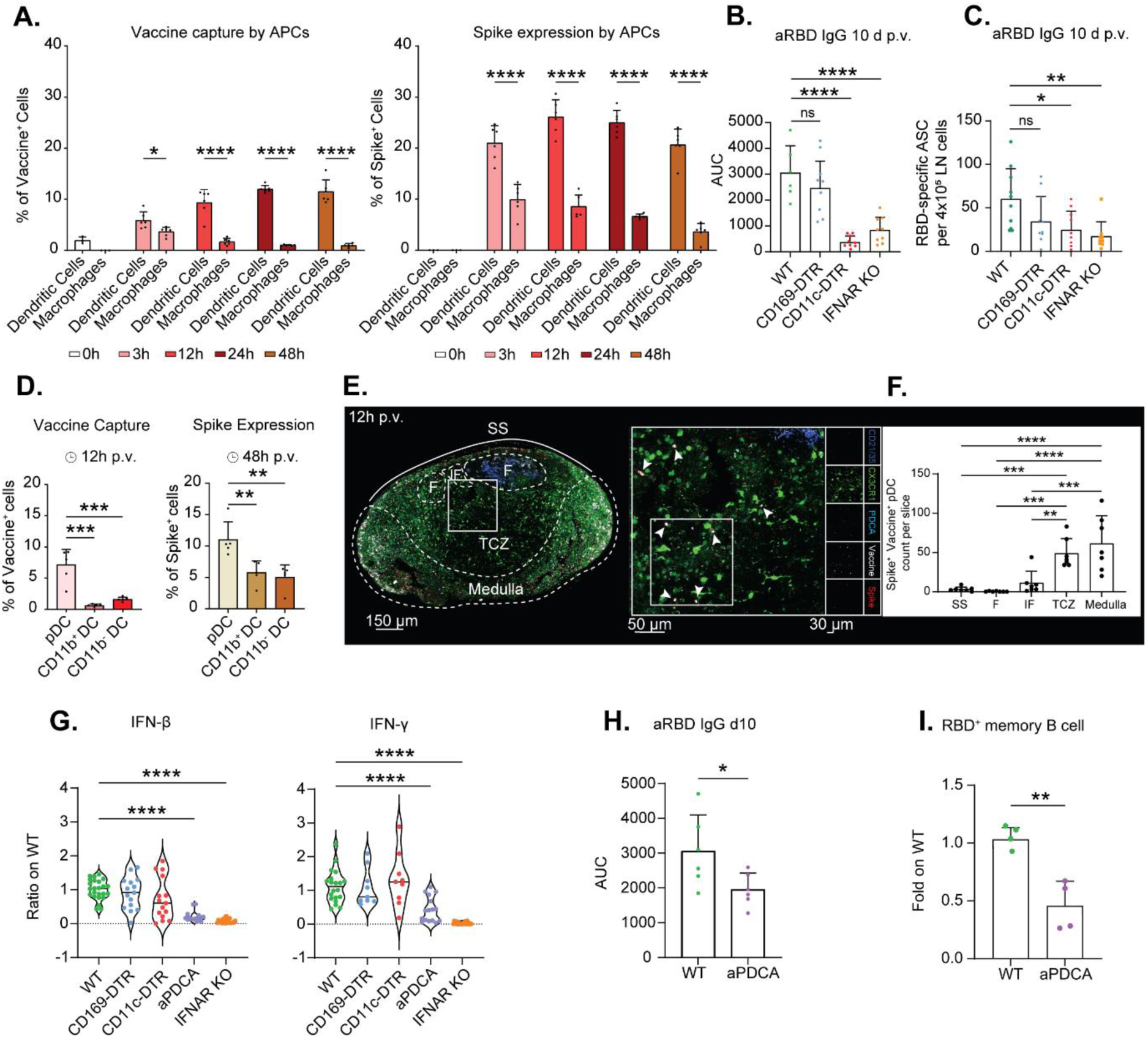
pDCs are essential to facilitate the humoral response following BNT162b2 vaccination. **(A)** Histogram showing the frequency of immune cells positive for the BNT162b2 vaccine (left) and the coronavirus spike protein (right) in the pLN at different time points p.v. (n=3 for 0h group, n=6 for all the other groups). **(B)** Area under the curve (AUC) indicating anti-SARS-CoV-2 RBD IgG titer for WT, CD169-DTR, CD11c-DTR, and IFNAR KO mice at day 10 p.v. (n=6 for WT group, n=9 for all the other groups. **(C)** Local anti-SARS-CoV-2 RBD IgG response measured by ELISPOT in WT, CD169-DTR, CD11c-DTR, and IFNAR KO mice at day 10 p.v. (n=10). **(D)** Histogram showing the capture of BNT162b2 vaccine at 12 h p.v. (left) and the expression of coronavirus spike protein at 48 h p.v. (right) by pDCs, CD11b^+^ DCs, and CD11b^−^ DCs in the pLN. (n=5). **(E)** Confocal micrograph showing a transverse section of a pLN (left) and magnifications of the T cell zone (TCZ, right) indicating vaccine capture and spike expression in PDCA^+^ pDCs at 12 h p.v. Colors indicate CD21/35^+^ follicles (blue), CX3XR1^+^ cells (green), PDCA^+^ pDCs (cyan), BNT162b2 vaccine (white), and spike protein (red). F stands for B cell follicle. **(F)** Quantification of spike^+^ pDCs in the different areas of the pLN at 12 h p.v. **(G)** Violin plots showing the ratio of IFN-β (left) and IFN-γ (right) expression in the lymph of CD169-DTR, CD11c-DTR, aPDCA, and IFNAR KO mice compared to WT mice at 12 h p.v. (Left graph: n=20 for WT group, n=15 for CD169-DTR, CD11c-DTR and aPDCA groups, n=24 for IFNAR KO group; Right graph: n=20 for WT group, n=9 for CD169-DTR, CD11c-DTR/GFP groups, n=16 for aPDCA group, n=24 for IFNAR KO group). **(H)** AUC for anti-SARS-CoV-2 RBD IgG in WT and aPDCA-treated mice at day 10 p.v. (n=6). **(I)** Flow cytometric analysis showing the ratio of SARS-CoV-2 RBD specific memory B cells in aPDCA mice compared to WT at day 10 p.v. (n=4). **(B-C, H)** The graph is representative of three independent experiments. Data are presented as mean ± SD. Two-way ANOVA **(A)** and One-way ANOVA **(B-G)** followed by Bonferroni correction for multiple comparisons. Student’s t-tests **(H-L)** (* p < 0.05, ** p < 0.01, *** p < 0.001 **** p<0.0001).

### Single-cell transcriptomic analysis predicts an interaction between pDCs and CD8^+^ T cells

To explore potential cell-cell interaction involving pDCs, we analyzed single-cell transcriptomic data from dLNs following BNT162b2 vaccination^28^. With the CellChat analysis tool^30^, we quantitatively characterized the intracellular communication network. This analysis was performed at 0 h, 24 h, and 7 days p.v.. Using dimensionality reduction via uniform manifold approximation and projection (UMAP) and graph-based clustering, we identified 18 cell clusters encompassing all major innate and adaptive immune cell types (**Fig. 3A**). Subsequent analysis enabled visualization of intercellular communication dynamics over time (**Fig. 3B**). Our investigation focused on the direct interaction of pDCs with B cells and their potential role in T-dependent antibody response, mainly through interactions with T cells or cDCs. To address this, we evaluated the probability of communication between pDCs and B cells, T cells, or cDCs over time. Notably, we identified a higher communication probability between pDCs and T cells compared to B cells or cDCs (**Fig. 3C**). Ligand-receptor (L-R) analysis further identified unique interactions at 24 h p.v., including complexes such as H2-q10 – CD8a, H2-q10 – CD8b1, H2-t24 – CD8a, H2-t24 – CD8b1, Icosl – CD28, Icosl – Icos. These findings suggest a communication between pDCs and CD8^+^ T cells (**Fig. 3D**).

**Fig. 3.**
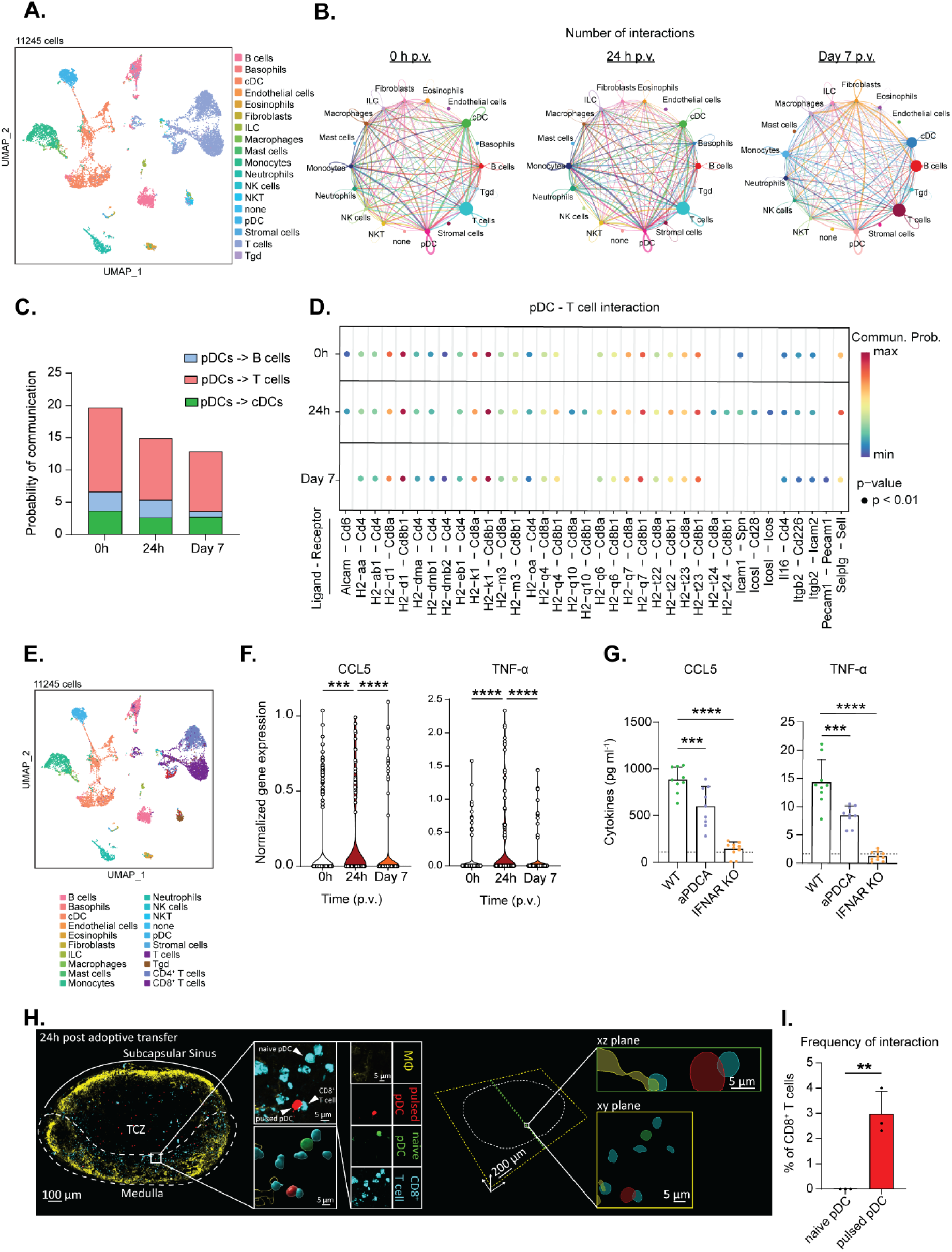
Cell-cell interaction analysis predicts communication between pDCs and CD8+ T cells following vaccination. **(A)** UMAP of cell types clustered by single-cell transcriptional analysis of 11245 cells after quality control. ILC NK stands for innate lymphoid cells and natural killer cells, respectively. **(B)** Circle plot showing the intercellular communication between the major immune cell types in the dLN at 0 h, 24 h, and 7 days p.v. **(C)** Histogram showing the probability of communication between pDCs with T cells, B cells, or cDCs at 0 h, 24 h, and 7 days p.v. **(D)** Comparison of the significant ligand-receptor pairs between pDCs and T cells in the dLN p.v. Dot color reflects communication probabilities, and dot size represents computed p-values. Space indicates that the communication probability is zero. p-values are calculated from one-sided permutation test. **(E)** UMAP of cell types clustered by single-cell transcriptional analysis of 11245 cells after quality control. **(F)** Truncated violin plots showing the gene expression of CCL5 (left) and TNF-α (right) expressed by pDCs at single-cell levels at 0 h, 24 h, and 7 days p.v. Each dot represents a single cell. **(G)** Histogram showing the expression levels of CCL5 (left) and TNF-α (right) in the LN following administration of the BNT162b2 vaccine in WT animals, in a group treated with aPDCA and in IFNAR KO mice at 12 h p.v. (n=9 for WT and aPDCA group, n=10 for IFNAR KO group). The negative control group (PBS) is indicated with a dashed line. **(H)** A representative confocal micrograph shows a transversal section of the pLN (left) and a magnification view showing the interaction between a pulsed pDCs and a CD8^+^ T cells (center). A representative mask shows the overlapping of the pulsed pDCs and CD8^+^ T cells signals (Right). Colors indicate CD169^+^ macrophages (MΦ) (yellow), vaccine-pulsed pDCs (red), naïve pDCs (green), and CD8^+^ T cells (cyan). **(I)** Histogram showing the frequency of CD8^+^ T cells interacting with naïve pDCs or in vitro pulsed pDCs 24 h post adoptive transfer. Data are presented as mean ± SD. One-way ANOVA followed by Bonferroni correction for multiple comparisons **(F-G)**. Student’s t-tests **(I)** (* p < 0.05, ** p < 0.01, *** p < 0.001 **** p<0.0001).

To validate the L-R interactions, we re-annotated cell clusters to differentiate between CD4^+^ and CD8^+^ T cells and assessed the expression levels of the respective ligand-receptor (L-R) genes (**Fig 3E**). At 24 h p.v., we observed increased Icosl, H2-q10, and H2-t24 expression in pDCs compared to baseline (0 h) and day 7. Additionally, pDCs activation was further corroborated by the upregulation of CD69 and the mRNA sensing gene Ifih1 (**Suppl. Fig. 3A**). Concurrently, we found that CD8^+^ T cells displayed an increased expression of Icos and CD28, as well as a significant expression of the activation markers CD69 and Gzmb at 24 h p.v. (**Suppl. Fig. 3B**). To support the predicted CD8^+^ T cell - pDC interaction, we also analyzed the expression of CCL5 and TNF-α, confirming an upregulated gene expression at 24 h p.v. compared to both 0 h and day 7. To confirm these results, we evaluated the expression of CCL5 and TNF-α in vaccinated aPDCA-treated and *IFNAR KO* mice compared to WT mice at 12 h p.v. As expected, depletion of pDCs or loss of IFNAR signaling markedly reduced the levels of CCL5 and TNF-α compared to the control groups (**Fig. 3F, G**). To support the predicted interactions previously identified, we performed adoptive transfer experiments with fluorescently labeled naïve CD8^+^ T cells, naïve pDCs, and *in vitro* pulsed pDCs in C57BL/6 mice. After 24 h, the pLNs were collected and analyzed by confocal microscopy. The results revealed a significant increase in the interactions between vaccine-pulsed pDCs and CD8^+^ T cells. In contrast, naïve pDCs failed to establish such interactions (**Fig. 3H, I**). These findings suggest a potential role of spike⁺ pDCs in directly engaging and activating CD8⁺ T cells within the pLN, thereby facilitating their functional differentiation post-vaccination.

### Spike^+^ pDCs engage with CD8^+^ T cells, enhancing their activation in the pLN

To investigate the phenotypic changes in the CD8^+^ T cell population following vaccination, we performed cluster analysis using the Phenograph algorithm, which identified 10 distinct CD8^+^ T cell subsets (**Fig. 4A**), exhibiting dynamic changes over time (**Fig. 4B; Suppl. Fig. 4A**). A heatmap displaying the normalized marker expression of CD44, CD62L, Ki-67, and MHC I across clusters allowed us to determine cluster identities (**Fig. 4C**). Quantitative analysis of CD8^+^ T cells frequencies revealed a significant decline in naïve CD8^+^ T cells at 24 h p.v., coinciding with an increase in active double negative (DN) and active naïve CD8^+^ T cells at the same time point (**Fig 4D**). By day 7 p.v., this transition culminated in a notable expansion of proliferating naïve CD8^+^ T cells (**Fig 4D, Suppl. Fig. 4B**). Conversely, other clusters identified by the UMAP analysis did not exhibit statistically significant differences over time (**Suppl. Fig. 4C**). Overall, these results describe the activated state of CD8^+^ T cells post-vaccination.

**Fig. 4.**
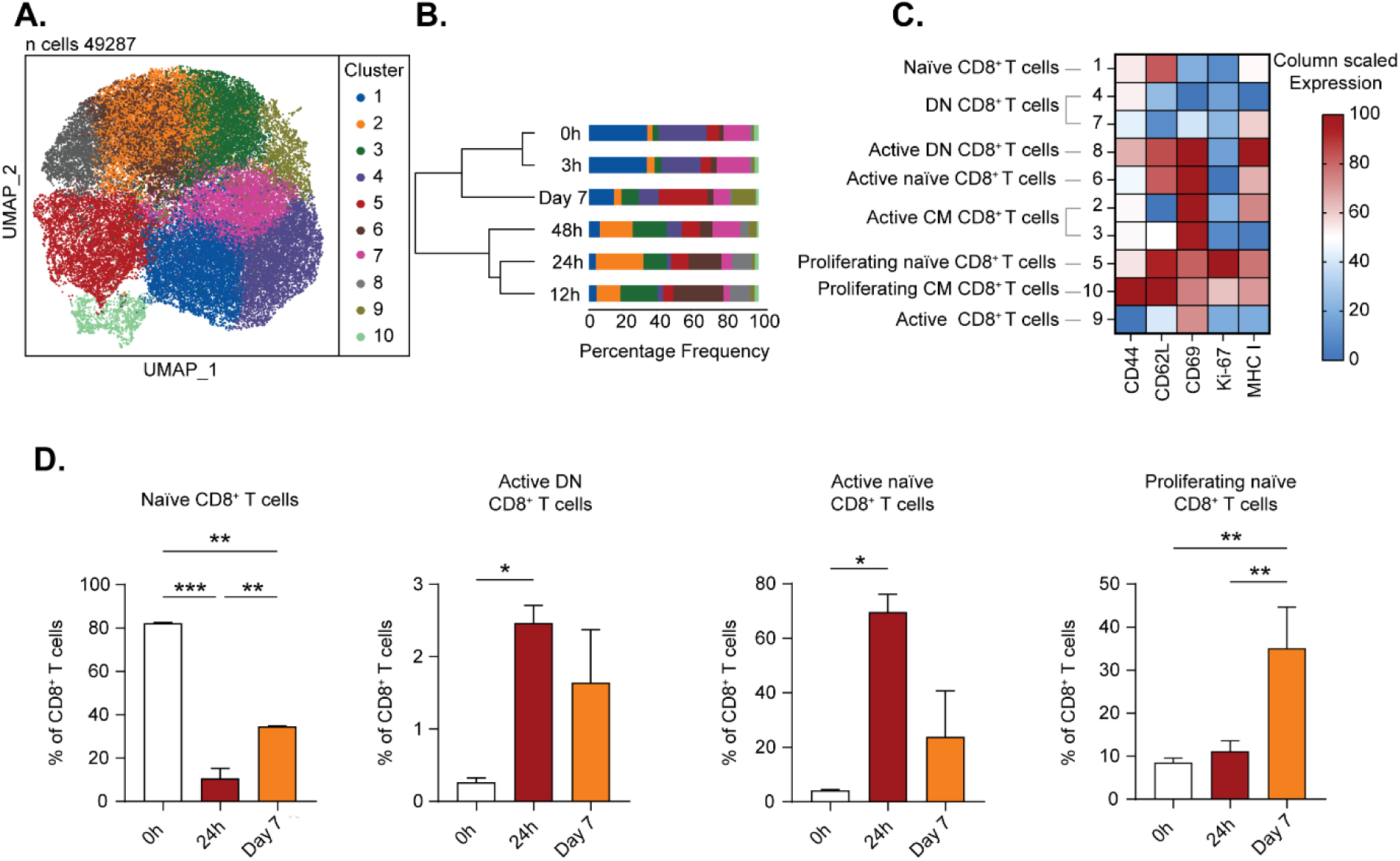
T cell activation induced by BNT162b2 vaccine in the pLN. **(A)** UMAP plot showing the clusters of CD8^+^ T cell obtained from flow cytometrical analysis of 49287 cells p.v. **(B)** Stacked-bar graph showing the percentages of CD8^+^ T cells Phenograph clusters. **(C)** Heatmap showing the column-scaled expression of markers (columns) in discrete CD8^+^ T cell Phenograph clusters (rows). **(D)** Histogram showing the main CD8^+^ T cell subsets at 0 h, 24 h, or 7 days p.v. in the pLN (n=3 for each group. Data are presented as mean ± SD. One-way ANOVA followed by Bonferroni correction for multiple comparisons (* p < 0.05, ** p < 0.01, *** p < 0.001 **** p<0.0001).

### Spike^+^ pDCs are essential for antigen-specific memory T cell formation following vaccination

Several studies have confirmed the induction of antigen-specific T cells after BNT162b2 vaccination^31,32^. Additionally, type I IFNs have been shown to directly stimulate CD8^+^ T cells, promoting clonal expansion and memory formation^31^. Based on these findings, we investigated whether pDCs depletion impairs the T-cell response to vaccination.

To address this point, we analyzed the lymphocyte infiltration in the lungs of vaccinated WT, aPDCA-treated, and *IFNAR KO* mice (**Fig. 5A**). Both aPDCA-treated and *IFNAR* KO mice exhibited a marked reduction in lymphocyte infiltration, comparable to non-vaccinated controls (**Fig. 5B**). Further analysis of T and B cell ratios revealed that only vaccinated WT mice, but not aPDCA-treated or *IFNAR KO* mice, showed T cell enrichment (**Fig. 5C**). Supporting our hypothesis, the depletion of pDCs or the absence of IFNAR resulted in a significant decrease in the overall memory CD4^+^ and CD8^+^ T cell populations (**Fig. 5D-E**). Notably, the spike-specific CD8^+^ T cell subset experienced the most pronounced reduction, underscoring the critical role of pDCs in promoting T cell memory formation (**Fig. 5F**).

**Fig. 5.**
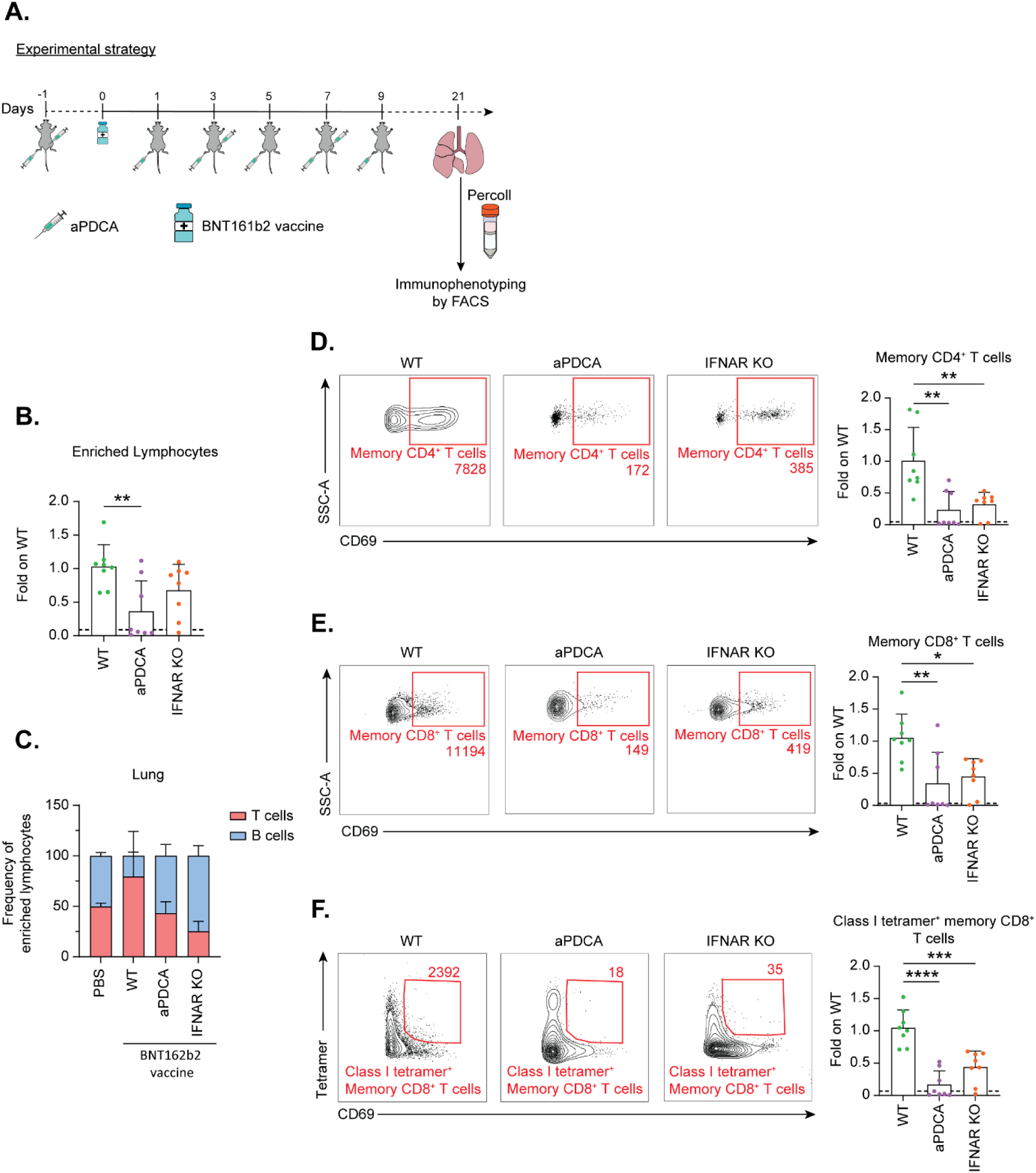
Spike^+^ pDCs are important for BNT162b2 vaccine-induced memory CD8^+^ T cells formation. **(A)** Experimental strategy. **(B)** Histogram showing CD45^+^ cell counts in the Percoll-enriched lymphocyte fraction from the lungs of PBS vaccinated (dashed line) or BNT162b2 vaccinated WT, aPDCA-treated, and IFNAR KO mice, measured on day 21 p.v. **(C)** Stacked bar graph showing the frequency of B and T cells in the Percoll-enriched lymphocyte fraction from the lungs of PBS vaccinated or BNT162b2 vaccinated WT, aPDCA-treated, and IFNAR KO mice, measured on day 21 p.v. **(D)** Flow cytometry contour plot (above) from a representative experiment out of 3 and histogram (below) showing memory CD4^+^ T cells count in the Percoll-enriched lymphocyte fraction from the lungs of PBS vaccinated (dashed line) or BNT162b2 vaccinated WT, aPDCA and IFNAR KO mice, measured on day 21 p.v. **(E)** Flow cytometry contour plot (above) from a representative experiment out of 3 and histogram (below) showing memory CD8^+^ T cells count in the Percoll-enriched lymphocyte fraction from the lungs of PBS vaccinated (dashed line) or BNT162b2 vaccinated WT, aPDCA-treated and IFNAR KO mice, measured on day 21 p.v. **(F)** Flow cytometry contour plot (above) from a representative experiment out of 3 and histogram (below) showing Class I tetramer^+^ memory CD8^+^ T cells count in the Percoll-enriched lymphocyte fraction from the lungs of PBS vaccinated (dashed line) or BNT162b2 vaccinated WT, aPDCA-treated and IFNAR KO mice, measured on day 21 p.v. **(B-F)** The experimental groups were made of n=3 for PBS, n=8 for all the other groups. Data are presented as mean ± SD. One-way ANOVA followed by Bonferroni correction for multiple comparisons (* p < 0.05, ** p < 0.01, *** p < 0.001 **** p<0.0001).

## Discussion

This study intended to elucidate the mechanisms by which the BNT162b2 vaccine elicits effective humoral and cellular immunity against SARS-CoV-2.

While previous studies described vaccine arrival in the LN ^33^, it remained unclear whether this transport was associated with migrating APCs or occurred via lymphatic drainage^27,34^. Here, we demonstrated that soon following injection, the BNT162b2 vaccine drains via the lymphatic system. Indeed, while the immune cells expressing the viral antigen require one to four days to migrate from the injection site to the dLN^35,36^, the inflammatory response is initiated in this organ already at three hours post-vaccination. This rapid activation highlights the role of LN-resident leukocytes in the early uptake and presentation of the antigen. However, this process might be influenced afterward by the migration of immune cells from the injection site. The impact of these events on the immune response to the vaccine needs to be further evaluated. Notably, the size and the surface composition of the drug carrier can significantly affect the pharmacokinetics and biodistribution properties of the associated small molecules^37^. For example, Cordeiro and colleagues showed how different sizes and surface properties of polymeric nanocapsules influence their ability to target specific regions of the LN, consequently modulating immune cell engagement^38^. Based on these studies, we could expect differences in the capture and distribution of other commercially available coronavirus vaccines in the dLN^39^. Indeed, we observed a preferential accumulation of the BNT162b2 vaccine in the interfollicular and medullary regions of the LN. This is most likely due to the convergence of a high number of medullary sinuses, increasing the draining surface, and the presence of different types of phagocytic populations^40^, responsible for the rapid capture of the vaccine and the expression of the spike protein. The distribution of the BNT162b2 vaccine was consistent with previous observations that identified the interfollicular region and the medulla as the key targets not only for inactivated vaccines (as the influenza type I vaccine)^17^, which are often considered highly effective natural vehicles for nucleic acid delivery, but also for polymeric nanoparticles^41,42^.

Consistent with previous studies highlighting the role of APCs during mRNA vaccination^28^, we observed that pDCs exhibited the highest vaccine uptake and spike protein expression following vaccination. Although the capacity of pDCs to capture nanoparticles has been demonstrated *in vitro*^43^, their involvement *in vivo,* especially in the context of vaccination, is still poorly understood. Brewitz and colleagues studied the localization of pDCs at steady state in the LN, finding a higher abundance of these cells in the interfollicular, paracortical, and medullary areas, preferentially associated with high endothelial venules. These authors also found that pDCs migrate towards the medulla and the T cell zone upon vaccination^44^. In our model, the vaccine accumulates in the same regions where pDCs are located, which could explain their higher capture compared to other immune cells. In addition, different studies have shown that viral RNA can be transferred between infected cells and pDCs through exosomes^45,46^. Therefore, the expression of the spike protein observed in the pDCs could be the result of the direct capture of the vaccine by the pDCs, but also the result of the transfer of the RNA or the proteins expressed by other cells located in the same area^44^.

By analyzing the contribution of pDCs in establishing a sustained adaptive immune response following vaccination, we found that the absence of this population significantly compromised the development of the adaptive response, affecting both the humoral and the cell-mediated response. Regarding the humoral response, it is not completely understood how B cells encounter their antigen. Different mechanisms have been proposed for antigen presentation to B cells upon infection or vaccination. in case of small antigens (<70 kDa) their capture occurs following a direct drainage via the conduit system located in the B follicles^47^. Conversely, larger antigens are captured by macrophages or cDCs in the subcapsular sinus and medullary regions and are transferred to the B cells^20^. Taking this into account we hypothesize that pDCs might relocate towards the B cell follicle and present the newly express antigen directly to the B cells or transfer it to the FDC network.

Furthermore, we, and others, have previously demonstrated the critical role of the type I IFN signaling in the development of a proper antibody response upon immunization^21,28,48^. In this work, we proposed an additional scenario in which the contribution of pDCs to the humoral response is associated with their role in the initiation of the type I IFN response that follows vaccination^18,29,48^. Here we identified pDCs as the primary source of the IFN-β in the dLN. Furthermore, Li and colleagues demonstrated that the mRNA vaccine is sensed by LN-resident cells by engaging the MDA5-IFNAR1 signalling pathway^28^. In addition, activated macrophages were shown to enhance type I IFN secretion by pDCs in the context of SARS-CoV-2 infection^49^. Supporting the involvement of pDCs in the sensing of the vaccine, we observed a strong upregulation of the related gene (*Ifih1*) following vaccination. This suggests that pDCs act as early sentinels for mRNA vaccines, integrating innate immune sensing with downstream adaptive immune activation. Taken together, these findings highlight a novel role of the pDCs in orchestrating early events that support the development of effective humoral immunity.

Along with B cells, type I IFN contributes to different immunological processes such as the T cell memory response against viruses^50^ or the migration of pDCs within the lymph node following viral infection^51,52^. In addition to that, a previous study showed that the MDA5-IFNAR1 signaling pathway is also critical for CD8^+^ T cell response induced by the BNT162b2 vaccine^28^. We extended these findings to show that spike^+^ pDCs were able to establish direct interactions with CD8^+^ T cells, suggesting their potential role in directly engaging and activating T cells within the dLN. Strengthening this hypothesis, we observed a significant loss of antigen-specific memory CD8^+^ T after immunization in mice lacking pDCs. Interestingly, similar pDCs - CD8^+^ T cell interactions have been described during viral infections^46,53^. For example, Mouriès and colleagues demonstrated the capacity of pDCs to cross-present exogenous antigen to naïve T cells, resulting in their activation and maturation^54^. This direct interaction provides mechanistic evidence that pDCs may function beyond their classical role in type I IFN production to act as unconventional APCs during vaccination.

The growing interest in pDCs in the cancer immunology field further supports their potential as targets for optimizing future vaccines ^42,55–58^. For example, Butkovich and colleagues showed how RNA-loaded nanoparticles can efficiently activate pDCs, improving the tumoral antigen presentation by cDCs to CD8^+^ T cells^43^.

In conclusion, our results provide novel insights into the *in vivo* mechanisms underlying the immune response to the BNT162b2 vaccine, with the discovery of a novel role of pDCs in mediating vaccine-induced immunity. Our findings contribute to a better understanding of nanoparticle-based vaccine-induced immune response and offer novel insights to improve vaccine efficacy in infectious and cancer immunotherapies.

## Materials and Methods

### Mice

C57BL/6J mice were bred in-house or purchased from Charles River Laboratories. B6.129P2(Cg)-Cx3cr1tm1Litt/J (CX3CR1-GFP) mice were initially purchased from Jackson Laboratories and subsequently bred in our animal facility at the Institute of Research in Biomedicine (IRB). B6.129(ICR)-Tg(CAG-ECFP)CK6Nagy/J (CK6-ECFP) mice were initially purchased from Charles River Laboratories and subsequently bred in our animal facility at the IRB. The following transgenic strains were used: CD169^DTR (59)^, CD11c^DTR/GFP (60)^, IFNAR^#x2212;/− (61)^. All animal experiments were conducted following the guidelines of the Swiss Federal Veterinary Service, and the protocols (animal permits TI 33/2021) were approved by the Animal Welfare Committee (Commissione Cantonale per gli Esperimenti sugli Animali) of the Cantonal Veterinary Office. Mice used for experiments included equal numbers of males and females, aged 6 to 12 weeks at the time of vaccination, in good health, and with no abnormal clinical signs. Mice were bred under specific pathogen-free (SPF) conditions in individually ventilated cages, with a controlled light-dark cycle (12:12), room temperature (20–24°C), and relative humidity (30%–70%). All the experiments were run in the basics of Biosafety Level 2 (BSL2) facility.

### Vaccination and injections

Discarded remnant material (Pfizer-BioNTech, BNT162b2) was stored at −80°C at the final concentration of 100 μg/ml. For fluorescence-based techniques, the vaccine was labeled with DiD (#V22889, Invitrogen) as previously described^62^. The final concentration was checked at the nanodrop, and the vaccine was diluted back to the initial concentration.

Pfizer-BioNTech mRNA vaccine (2μg in 10μl per mouse) was injected into each footpad of isoflurane-anesthetized mice (2% volume-to-volume, Baxter), and pLN was collected at different time points. For macrophage depletion, CD169^DTR^ mice were depleted 2 days before and 4 days after vaccination with 200 ng of diphtheria toxin (Sigma-Aldrich) intraperitoneally (i.p.) and subcutaneously (s.c.) in the footpad. For cDCs depletion, CD11c^DTR/GFP^ mice were depleted 2 days before and 4 days after vaccination by injecting 80 ng of diphtheria toxin (Sigma-Aldrich) s.c. in the footpad. For pDCs depletion, 400μg/mouse i.p. (day −1, +3, +7) and 100μg/footpad s.c. (day −1, +1, +3, +5, +7, +9) of αCD317 (clone 927, BioXCell) was administered to the mice.

### Two-photon intravital microscopy

The day before the surgery, mice were injected with 500ng αCD21/35 Pacific Blue s.c. in the footpad. On the day of the surgery, mice were injected with 500ng of αCD169 APC s.c. in the footpad. Mice were anesthetized with an i.p. injection of a cocktail containing ketamine (100mg/kg, Sintetica) and xylazine (10 mg/kg, Virbac). The mouse was positioned on a microscopic stage. The right foot was maintained and extended by fastening the fingers using a suture thread, and fasteners were attached to the hip. Finally, the pLN was exposed through an incision in the skin and by removing the overlying fat tissue. Mice were then injected s.c. in the footpad with 2μg of DiD-labelled BNT162b2 vaccine. The imaging of the pLN was performed on a customized up-right two-photon platform (TrimScope, LaVision BioTec) as previously described^63^. The objective used was a 10X, and two-photon micrographs were acquired every 30 sec for a total duration of 90 min. Analyses were performed using the Imaris 9.9.1 software (Oxford Instruments) and ImageJ^64^.

### Expression and purification of YFP-tagged RBD and RBD

Expression and purification of YFP-tagged RBD and RBD were performed with modifications to the methods described by Bierig et al.^65^. A DNA sequence encoding YFP-RBD-His was synthesized and cloned into the pcDNA3.1(+) vector by GenScript. Expi293F cells (#A14527, ThermoFisher) were cultured in Expi293 Expression Medium (#A1435101, ThermoFisher) in an orbital shaker at 120 rpm, 37°C, and 8% CO_2_. On the day of transfection, cells were adjusted to 3.0 × 10^6^ cells/mL and transiently transfected with 1 μg/mL pcDNA3.1(+) YFP-RBD-His plasmid DNA and 2.7 μg/μl DNA ExpiFectamine™ 293 Reagent (#A14524, ThermoFisher). Twenty-two hours post-transfection, ExpiFectamine™ enhancers 1 (#A14525, ThermoFisher) and 2 (#A14526, ThermoFisher) were added according to the manufacturer’s instructions. One week after transfection, the culture supernatant was supplemented with 10 mM imidazole (#I2399, Sigma-Aldrich), 300 mM NaCl (#S7653, Sigma-Aldrich), and 50 mM Tris (pH 7.5) (#T1503, Sigma-Aldrich), and centrifuged at 2000g for 30 min at 4°C. The clarified supernatant was loaded onto a HisTrap excel HP column (#17-3712-05, Cytiva) pre-equilibrated with binding buffer (50 mM Tris, pH 7.5, 10 mM imidazole, and 300 mM NaCl) using an AKTA purifier 10 FPLC system (Cytiva). The column was washed with five bed volumes of wash buffer (50 mM Tris, pH 7.5, 30 mM imidazole, and 300 mM NaCl), and proteins were eluted with three bed volumes of elution buffer (25 mM HEPES, pH 7.4, 300 mM NaCl, and 300 mM imidazole). Eluted peak fractions were pooled and dialyzed overnight in PBS (#D8537, Sigma-Aldrich). The dialyzed protein was concentrated to 2 mL using Amicon® Ultra Centrifugal filters (10 kDa cutoff, #UFC801024, Millipore). The concentrated protein was then injected into a Superdex 200 16/600 column (#28-9893-35, Cytiva) pre-equilibrated with PBS. Peak fractions were collected, concentrated using Amicon® Ultra Centrifugal filters, filter-sterilized with 0.2 micron Millex® syringe filters (#SLGV033RS, Millipore Sigma), and stored in small aliquots at −80°C. For the production of RBD protein without the YFP tag, all steps were identical except that after Ni-NTA purification, the dialyzed sample was treated with PreScission Protease (#27-0843-01, Cytiva) at a 1:100 ratio for 6 h at 4°C, and then passed through a GST resin (#17-5132-01, Cytiva) to remove the PreScission Protease. The flow-through was re-passed through the Ni-NTA resin to remove the cleaved YFP tag. As described above, the Ni-NTA eluted fractions were collected, concentrated, and injected into the Superdex 200 16/600 column for final purification.

### Tissue collection and preparation of single-cell suspension

Mice were sacrificed with CO_2,_ and organs were harvested. For LNs, after collection, they were disrupted with tweezers and filtered with a 40 µm cell strainer. For spleens, after collection, they were disrupted on a 40 µm cell strainer, and red blood cells (RBCs) were lysed using the Lysing Buffer (#555899, BD Biosciences). Then, CD8^+^ T cells or pDCs were isolated using the EasySep^TM^ mouse CD8^+^ T cell isolation kit (#19853, StemCell) and the mouse Plasmacytoid dendritic cell isolation kit (#130-107-093, Miltenyi). For lungs, mice were perfused with PBS, and lungs were processed as previously described^66^. RBCs were lysed, and lymphocytes were isolated from the lungs using a 40/80 gradient Percoll (VWR International), followed by centrifugation at 1250 rcf for 20 min without acceleration and break.

### Flow cytometry analysis of vaccine capture and spike expression

pLNs were processed as previously described. Single-cell suspensions were stained with antibodies listed in Table 1. Fc receptors were blocked with αCD16/32, and dead cell exclusion was performed using the Zombie Aqua^TM^ Fixable Viability Kit (Biolegend, San Diego, USA). Surface staining was carried out for 30 min, and vaccine capture was detected as DiD^+^ cells, while spike expression was detected using αSpike antibody. All operations were done at 4°C in the dark. B cells were identified as CD45^+^, CD3^−^, B220^+^, PDCA^−^. pDCs were identified as CD45^+^, CD3^−^, B220^+^, PDCA^+^. T cells were identified as CD45^+^, B220^−^, CD3^+^. In the CD45^+^, CD3^−^, B220^−^, all the downstream cells were gated. Neutrophils/monocytes were identified as GR1^+^. Medullary macrophages were identified as GR1^−^, F4/80^+^, and CD11b^+^. NK cells were identified as GR1^−^, F4/80^−^, and NK1.1^+^. DCs were identified as GR1^−^, F4/80^−^, NK1.1^−,^ and two major subpopulations were distinguished based on CD11b expression: CD11b^+^ cDCs and CD11b^−^ cDCs. Subcapsular sinus macrophages were identified as GR1^−^, F4/80^−^, NK1.1^−^, CD11c^−^, CD11b^+^. All samples were acquired on fluorescence-activated cell sorting (FACS) Symphony A3 flow cytometer (BD Biosciences, New Jersey, USA) and were analyzed with FlowJo software version 10.7.1 (FlowJo LLC, Ashland, Oregon).

### Flow cytometry analysis of memory B cells

SARS-CoV-2 RBD-YFP protein was produced in-house, as described above^65^. pLNs were processed as previously described. Single-cell suspensions were stained with antibodies listed in Table 1. Fc receptors were blocked with αCD16/32, and dead cell exclusion was performed using the Zombie Aqua^TM^ Fixable Viability Kit (Biolegend, San Diego, USA). Surface staining was carried out for 30 min. All operations were done at 4°C in the dark. RBD^+^ memory B cells were identified as CD45^+^, CD3^−^, B220^+^, CD43^−^, CD19^+^, CD1D^−^, IgG^+^, RBD^+^. All samples were acquired on fluorescence-activated cell sorting (FACS) Symphony A3 flow cytometer (BD Biosciences, New Jersey, USA) and were analyzed with FlowJo software version 10.7.1 (FlowJo LLC, Ashland, Oregon).

### Flow cytometry data analysis with Crusty

pLNs were processed as previously described. Samples were transferred in a 96-well plate, and they were stimulated with a mix of 50 μg/ml brefeldin (#B7651, Sigma-Aldrich), 20 ng/ml phorbol myristate acetate (PMA) (#10933, Sigma-Aldrich) for 3 h at 37°C. Single-cell suspensions were stained with antibodies listed in Table 1. Fc receptors were blocked with αCD16/32, and dead cell exclusion was performed using the Zombie Aqua^TM^ Fixable Viability Kit (Biolegend, San Diego, USA). Surface staining was carried out for 30 min at 4°C. Samples were then fixed and permeabilized in eBioscience^TM^ Foxp3/Transcription Factor Staining Buffer Set (#00-5523-00, ThermoFisher) and stained intranuclearly with αKi-67 PE in Fix/Perm buffer 30 min at 4°C. T cell subsets were distinguished based on the expression of CD4 and CD8 markers. For cluster discovery, the gates were downsampled to have 2900 CD8^+^ T cells per sample. Files were then exported as .csv files and imported to CRUSTY web platform. Clusters were determined by applying the PhenoGraph algorithm^67^. To study the identified clusters, CD8^+^ T cell subsets were gated as CD45^+^, B220^−^, CD3^+^, CD8^+^, in which the following subsets were identified: Proliferating Naïve CD8^+^ T cells Ki-67^+^, CD62L^+^, CD44^−^; Proliferating CM CD8^+^ T cells Ki-67^+^, CD62L^+^, CD44^+^; Naïve CD8^+^ T cells Ki-67^−^, CD69^−^, CD62L^+^, CD44^−/+^; DN CD8^+^ T cells CD8^+^, Ki-67^−^, CD69^−^, CD62L^−^, CD44^−/+^; Active Naïve CD8^+^ T cells Ki-67^−^, CD69^+^, CD62L^+^, CD44^−^; Active CM CD8^+^ T cells Ki-67^−^, CD69^+^, CD62L^+^, CD44^+^; Active DN CD8^+^ T cells Ki-67^−^, CD69., CD62L^−^, CD44^−/+^. All samples were acquired on fluorescence-activated cell sorting (FACS) Symphony A3 flow cytometer (BD Biosciences, New Jersey, USA) and were analyzed with FlowJo software version 10.7.1 (FlowJo LLC, Ashland, Oregon).

### Flow cytometry analysis of tissue-resident memory T cells in the lungs

Lungs were processed as described above. Then, lymphocytes were isolated with percoll gradient. Fc receptors were blocked with αCD16/32, and dead cell exclusion was performed using the Zombie Aqua^TM^ Fixable Viability Kit. Surface staining was carried out for 30 min at 4°C. T cells were identified as CD45^+^, B220^−^, CD3^+^. Then CD4^+^ and CD8^+^ T cells were identified, and tissue-resident cells were gated as CD69^+^. To conclude, SARS-CoV-2 specific memory T cells were identified as CD45^+^, B220^−^, CD3^+^, CD8^+^, CD69^+^, Class I tetramer^+^. All samples were acquired on fluorescence-activated cell sorting (FACS) Symphony A3 flow cytometer (BD Biosciences, New Jersey, USA) and were analyzed with FlowJo software version 10.7.1 (FlowJo LLC, Ashland, Oregon).

### Multiplex assay

pLNs were collected at different time points after vaccination and carefully disrupted in 75μL of cold PBS^#x2212;/−^to avoid cell rupture. The suspension was centrifuged at 1,500 rpm for 5 min, and the supernatant was collected and stored at −80°C until the usage. According to the manufacturer’s instructions, the concentration of cytokines in the lymph was determined by LEGENDPlex assays (#740150, Mouse Inflammation Panel (13-plex); #740622, Mouse Anti-Virus Response Panel (13-plex); Biolegend). Samples were analyzed using a Symphony A3 flow cytometer (BD Biosciences, New Jersey, USA), and data were analyzed using LEGENDPlex software (BioLegend).

### ELISA

SARS-CoV-2 RBD protein was produced in-house, as described above. Blood was collected from the tail vein in capillary blood collection system tubes (GK 150 SE Gel 200 µl, Kabe Labortechnik) on day 7 and day 10 p.v. Blood was then centrifuged at 3,000g for 10 min RT. The serum was aliquot into 1.5 ml Eppendorf tubes and stored at −80°C until the usage. The experiment was then performed as previously described. Half-well Nunc ELISA plates were coated with 5μg/ml SARS-CoV-2 RBD protein in PBS^#x2212;/−^ buffer and incubated overnight at 4°C. The experiment was then performed as previously described^62^.

### ELISpot

For enzyme-linked immunosorbent spot assay (ELISpot), on day 10 p.i., popliteal LNs were removed aseptically, disrupted, and passed through a 40-μm cell strainer. 4 ×10^5^ cells were plated on SARS-CoV-2 RBD-coated (5 μg/ml) filter plates (MultiScreen HTS, Merck Millipore) and incubated overnight at 37°C. For detection, a biotin-conjugated anti-IgG or anti-IgM was used, followed by streptavidin-conjugated horseradish peroxidase (HRP). A developing solution consisting of 200 μL 3-amino-9-ethylcarbazole (AEC) solution (Sigma-Aldrich) in 9 mL sodium acetate buffer containing 4 μL 30% H_2_O_2_ was subsequently added. Spots were read on a CTL ELISPOT reader using ImmunoSpot 5.1 software (Cellular Technology).

### *In vitro* pDCs pulsing with DiD-labelled BNT162b2 vaccine

1μg/ml DiD-labelled BNT162b2 vaccine was incubated in complete RPMI 1640 (#31870-025, Gibco) supplemented with 10% heat-inactivated fetal bovine serum (FBS) (#10270-106, Gibco), 50 units/ml penicillin + 50 μg/ml streptomycin (#15070-063, Gibco), 1% glutamax (#35050-038,Gibco), 1% sodium pyruvate (Gibco), 1% nonessential amino acids (#11140-035, Gibco) for 60 min at 37°C with 5% CO^2^ to allow the formation of the protein corona. pDCs were isolated as previously described. 3 × 10^5^ cells/ml pulsed pDCs or naïve pDCs were plated in a Petri dish with 1μg/ml DiD-labelled BNT162b2 vaccine or complete RPMI alone for 3 hours at 37°C.

### Immunofluorescence

For mouse microscopy experiments, pLNs were harvested at different time points and fixed in 4% paraformaldehyde (PFA; Merck-Millipore) for 12 h at 4° and then embedded in 4% Low Gelling Temperature Agarose (Sigma-Aldrich). Sequential sections of 40 μm were cut with a vibratome (VT1200S, Leica Microsystems) and stained in a blocking buffer composed of PBS supplemented with calcium and magnesium (PBS+/+), TritonX100 (VWR) 0.1%, BSA 5% (VWR), and fluorescently labeled antibodies at the appropriate concentration. After overnight incubation at RT, samples were washed in PBS+/+ with 0.05% Tween 20 (Sigma-Aldrich), followed by washes with PBS-/-, and mounted on glass slides. To study the vaccine uptake and Spike expression, CX3CR1-GFP mice were immunized as described in the “Vaccination and injections” section, and slices were stained for CD21/35 and spike. pLN regions were manually identified based on CX3CR1 and CD21/35 expression. To study pDC-T cell interaction, pDCs and CD8^+^ T cells were isolated as reported in the “Tissue collection and preparation of single-cell suspension” section from WT and CK6-ECFP mice, respectively. pDCs were then stimulated and pulsed as previously described, and naïve and pulsed pDCs were labeled with 1 μM CellTrace^TM^ CFSE Cell Proliferation kit (# C34554, ThermoFisher) and 1μM CellTracker^TM^ Deep Red (# C34565, ThermoFisher) respectively. 3 × 10^6^ naïve and pulsed pDCs and 4×10^6^ CD8^+^ T cells were injected intravenously in the mice. pLN slices were stained for CD169. Immunofluorescence confocal images were acquired using a Leica TCS SP5 confocal microscope (Leica Microsystems), and images were analyzed using ImageJ^64^ and Imaris 9.7.2 Cell Imaging Software (Oxford Instruments). The antibodies used are listed in Table 1.

### scRNA-seq data analysis

Public data from the study were retrieved from GEO (accession number GSE179131) and analyzed using Seurat (v5.1.0)^68^. Cells with fewer than 200 or more than 5000 genes or mitochondrial RNA content exceeding 8% were excluded from the analysis. Genes expressed in very few cells (less than 3) were discarded. Raw count data were log normalized with a scaling factor of 10000. Subsequently, the most variable features were identified using the variance stabilizing transformation method, selecting the top 2,000 features based on their variability across the dataset. Finally, the data were scaled to mean center and unit variance for each feature, utilizing all genes in the dataset. We utilized Seurat’s anchor-based integration method on the dataset. Cell types were annotated with the ImmGen database.

### Cell communication analysis

Single-cell RNA sequencing (scRNA-Seq) data from mouse samples were processed using Seurat (v4.0) in R^69^. Raw sequencing reads were preprocessed as described in Virgilio T. et al. ^70^. Briefly, Dimensionality reduction was performed using principal component analysis (PCA), and significant principal components (PCs) were selected based on an elbow plot and JackStraw analysis. The top PCs were used to construct a shared nearest neighbor (SNN) graph, and clustering was performed using the FindClusters function with the Louvain algorithm. Uniform Manifold Approximation and Projection (UMAP) was applied for visualization. Cell types were annotated based on the expression of canonical marker genes and verified by comparison with publicly available single-cell datasets and literature. To infer intercellular communication between cell populations of interest, we employed CellChat (v1.5.0)^30^, a computational framework that predicts ligand-receptor interactions from scRNA-Seq data. The normalized gene expression matrix from Seurat was imported into CellChat using the createCellChat function. The CellChatDB.mouse database was used to identify potential ligand-receptor pairs specific to the mouse model. The communication probability was computed using a probabilistic model, and key signaling pathways were inferred using computeCommunProb and computeCommunProbPathway functions. The strength and directionality of interactions between cell populations were visualized using netVisual_circle, netVisual_bubble, and netAnalysis_contribution functions. The top ligand-receptor interactions were further examined in the context of known cellular functions and disease-related pathways. Statistical significance for differential ligand-receptor interactions was assessed using permutation-based significance testing implemented in CellChat. Comparisons between groups were performed using the Wilcoxon rank-sum test, and p-values were adjusted for multiple comparisons using the Benjamini-Hochberg method. A significance threshold of p < 0.05 was applied. All scripts and processed data this study uses are available upon reasonable request. Raw sequencing data have been deposited in Gene Expression Omnibus (GEO) under accession number Gene-at GSE212227.

### Statistical analysis

All statistical analyses were performed with Prism (GraphPad Software v10.4.1). For two-group comparisons, p values were determined using Student’s t-tests (two-tailed). For more than two groups, a comparison of one-way or two-way ANOVAs followed by Bonferroni correction was applied. Differences between groups were considered significant for p values < 0.05.

## Acknowledgments

The present research was funded by the Leonardo Foundation and the Swiss National Science Foundation (SNSF 204636). We thank the National Institutes of Health tetramer core facility for providing the spike-specific tetramer for CD8+ T cell detection.

## Author contributions

**C. Pizzichetti:** Conceptualization, data curation, formal analysis, investigation, visualization, methodology, writing–original draft, project administration, writing– review and editing. **I. Latino:** Conceptualization, data curation, formal analysis, funding acquisition, investigation, visualization, methodology, writing–original draft, project administration, writing– review and editing. **T. Virgilio**: Investigation, methodology. **A. Capucetti**: Investigation, methodology. **K. Chahine**: Investigation. **L. Cascione**: Data curation, formal analysis, visualization, methodology. **S. Moro**: Data curation, formal analysis, visualization, methodology. **M. Brügger**: Resources, investigation, methodology. **N. Kozarak:** Resources, investigation, methodology. **A. Pulfer**: Investigation, methodology, formal analysis. **L. Renner**: investigation. **R. S. Thakur**: Investigation, methodology. **C. Benarafa**: Resources, investigation, methodology. **D. F. Legler**: Conceptualization, discussion, supervision. **S. F. Gonzalez**: Conceptualization, resources, data curation, supervision, funding acquisition, writing–original draft, project administration, writing–review and editing.

## Supplementary Table

**Table 1.** Antibody list.

| Antibody | Fluorophore | Company | Cat. Num | Clone |
| --- | --- | --- | --- | --- |
| F4/80 | FITC | Biolegend | 123120 | BM8 |
| CD86 | PercCP | Biolegend | 105026 | GL-1 |
| Gr1 | PE-CF594 | BD<br>Bioscience | 562700 | 1A8 |
| CD11c | BV711 | Biolegend | 117349 | N418 |
| CD11b | BUV395 | BD<br>Bioscience | 563553 | M1/70 |
| B220 | BUV496 | BD<br>Bioscience | 612950 | RA3-6B2 |
| CD45 | BUV563 | BD<br>Bioscience | 612924 | 30-F11 |
| NK1.1 | BUV615 | BD<br>Bioscience | 751111 | PK136 |
| CD3 | PECY7 | Biolegend | 100220 | 17A2 |
| CD21/35 | Pacific Blue | Biolegend | 123414 | 7 E9 |
| CD11c | BV711 | Biolegend | 117349 | N418 |
| CD169 | AF647 | Biolegend | 142408 | 3D6.112 |
| CD16/32 | Purified | Biolegend | 101302 | 93 |
| Zombie aqua fixable viability kit |  | Biolegend | 423101 |  |
| CD19 | BV605 | Biolegend | 115540 | 6D5 |
| IgG | PE-CF594 | BD<br>Bioscience | 562538 | G18-145 |
| CD4 | BV786 | Biolegend | 100453 | GK1.5 |
| CD3 | PerCP-eFluor710 | ThermoFisher | 46-0032-80 | 17A2 |
| CD3 | BV711 | Biolegend | 100241 | 17A2 |
| PDCA | BV605 | Biolegend | 127025 | 927 |
| SARS-CoV-2 Spike S1 Subunit | BV750 | Biotechne | FAB105805S | 1035226 |
| SARS-CoV-2 Spike S1 Subunit | DyLight 594 | Biotechne | FAB105805M | 1035226 |
| PDCA | PE | Biolegend | 127009 | 927 |
| CD1d | BV421 | Biolegend | 123517 | 1B1 |
| CD43 | PE | BD Bioscience | 5532271 | S7 |
| IgM | PE-Cy7 | Biolegend | 406513 | RMM-1 |
| CD8 | BV650 | Biolegend | 100742 | 53-6.7 |
| CD44 | APC | Biolegend | 103017 | IM7 |
| CD62L | PercP | Biolegend | 104430 | MEL-14 |
| CD69 | FITC | Biolegend | 104506 | H1.2F3 |
| Ki-67 | PE | ThermoFisher | 12-5698-80 | SolA15 |
| MHC I | BV421 | Invitrogen | 48_9598_82 | AF6_88.5.5.3 |
| B220 | APC-H7 | Miltenyi | 130-110-849 | REA775 |
| CD103 | PercP Cy5.5 | Biolegend | 121416 | 2.00E+07 |

---

**Suppl. Fig. 1.**
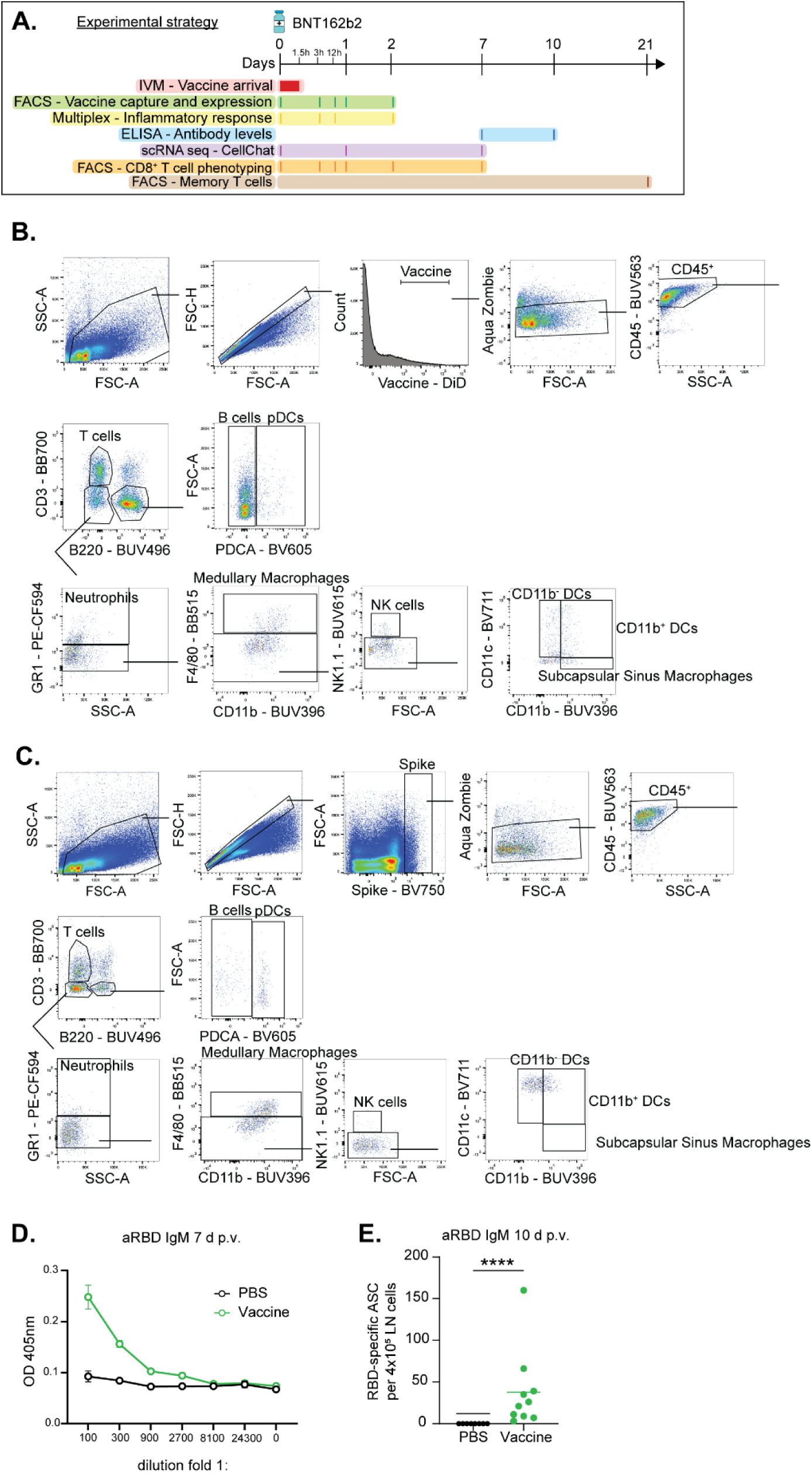
BNT162b2 vaccine induces a robust antibody response in mice. **(A)** Experimental strategy. IVM, Intravital microscopy; FACS, fluorescence-activated cell sorting; ELISA, enzyme-linked immunosorbent assay. **(B)** A gating strategy was used to identify different populations of bnt162B2 vaccine^+^ leukocytes. B cells and pDCs were identified as CD45^+^/CD3^−^/B220^+^ cells, and they were distinguished by the expression of PDCA. Meanwhile, T cells were gated CD45^+^/B220^−^/CD3^+^ cells. Neutrophils were identified in the gate of CD45^+^/CD3^−^/B220^−^ as positive for GR1. Within the negative population, Medullary macrophages were gated as F4/80^+^/CD11b^+^. NK cells were identified as CD45^+^/CD3^−^/B220^−^/GR1^−^/F4/80^−^/NK1.1^+^. To conclude, in the CD45^+^/CD3^−^/B220^−^/GR1^−^/F4/80^−^ /NK1.1^−^ it was possible to identify CD11b^−^ DCs as CD11c^+^/CD11b^−^, CD11b^+^ DCs as CD11c^+^/CD11b^+^, and medullary macrophages as CD11c^−^/CD11b^+^. **(C)** Systemic anti-SARS-CoV-2 RBD IgM titers were measured by ELISA on day 7 p.v. **(D)** Local anti-SARS-CoV-2 RBD IgM response measured by ELISPOT on day 10 p.v. Data are presented as mean ± SD. Mann-Whitney U test. (* p < 0.05, ** p < 0.01, *** p < 0.001 **** p<0.0001).

**Suppl. Fig. 2.**
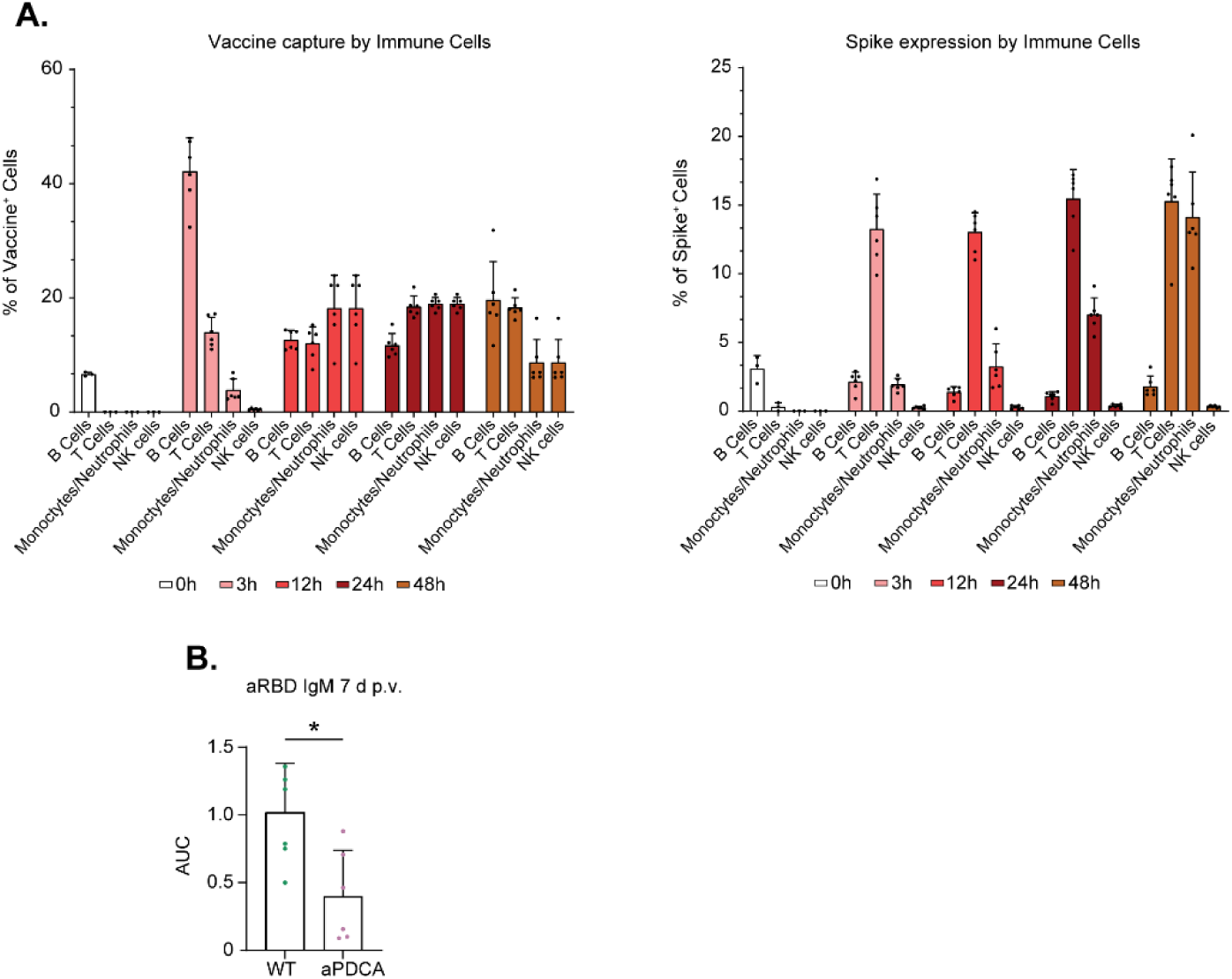
pLN activation following BNT162b2 vaccination. **(A)** Histogram showing the frequency of DiD-labeled BNT162b2 vaccine^+^ (left) and coronavirus spike protein^+^ (right) immune cells in the pLN at different time points p.v. (n=3 for 0 group, n=6 for all the other groups). **(B)** AUC for systemic anti-SARS-CoV-2 RBD IgM for WT and aPDCA-treated mice at day 7 p.v. (n=6 for each group). The graph is representative of two independent experiments. Data are presented as mean ± SD. One-way ANOVA followed by Bonferroni correction for multiple comparisons. Student’s t-tests (* p < 0.05, ** p < 0.01, *** p < 0.001 **** p<0.0001).

**Suppl. Fig. 3.**
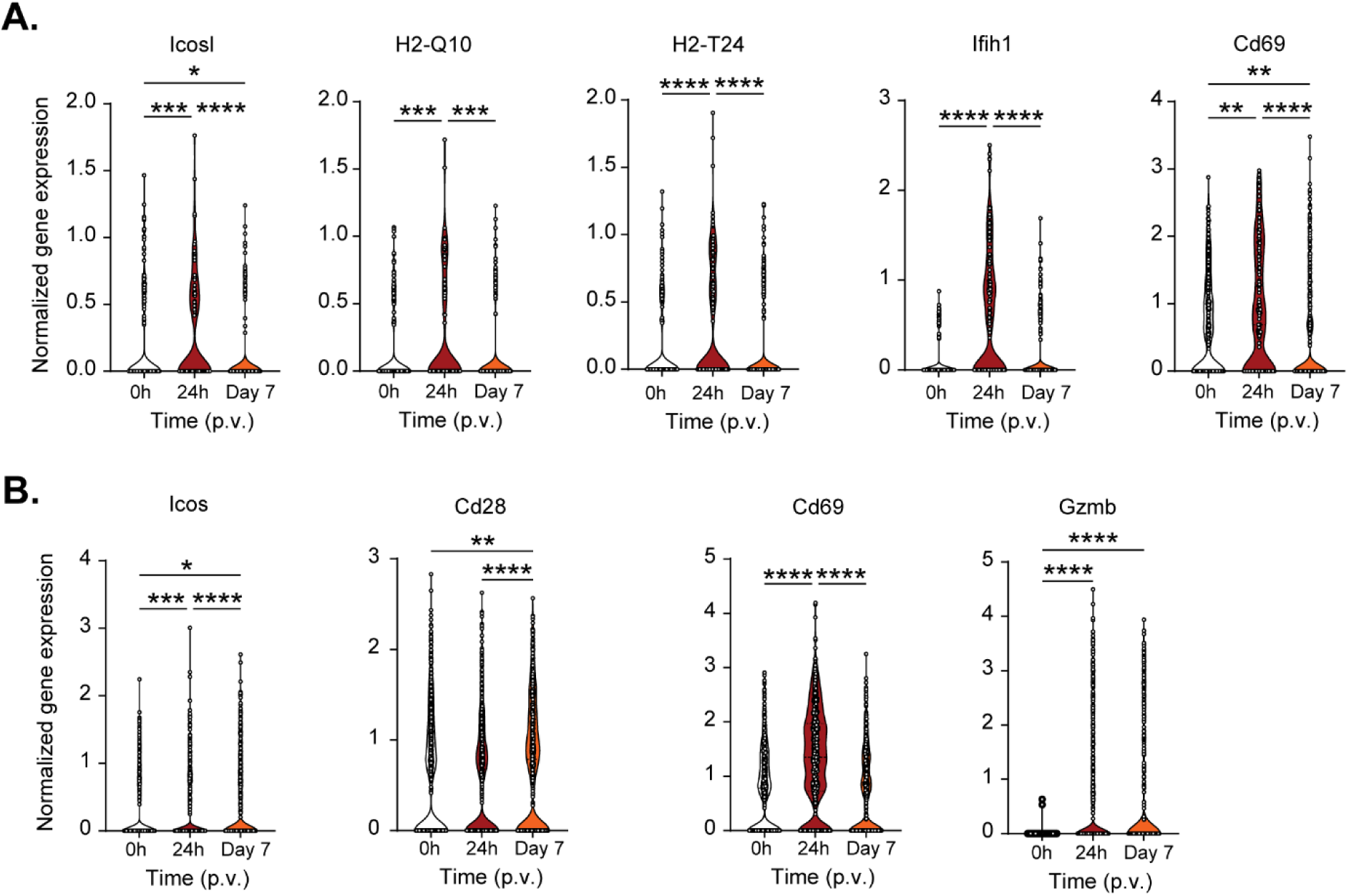
pDCs and CD8^+^ T cells gene expression in the pLN of vaccinated mice. **(A)** Truncated violin plots showing gene expression of pDCs on the single-cell level at 0 h, 24 h, and 7 days p.v. Each dot represents a single cell. **(B)** Truncated violin plots showing gene expression of CD8^+^ T cells on the single-cell level at 0 h, 24 h, and 7 days p.v. Each dot represents a single cell. Data are presented as median ± SD. One-way ANOVA followed by Bonferroni correction for multiple comparisons (* p < 0.05, ** p < 0.01, *** p < 0.001 **** p<0.0001).

**Suppl.Fig. 4.**
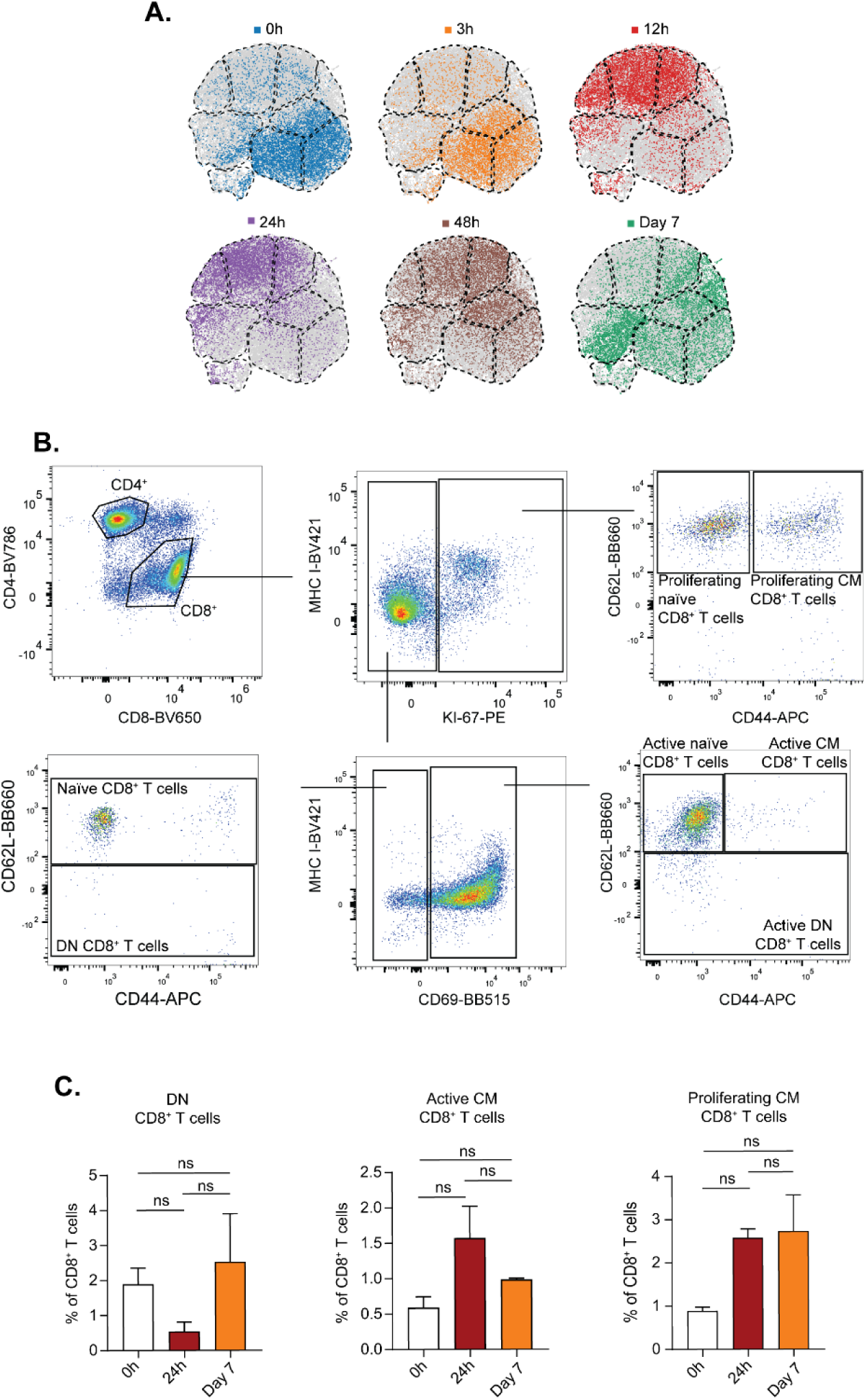
T cell phenotype in the pLN of vaccinated mice. **(A)** plots showing single CD8^+^ T cells from specific sets of samples. The grey background refers to the total CD8^+^ T cells in the dataset. **(B)** The gating strategy was used to identify different subsets of CD8^+^ T cells, which were gated as single/viable/CD45^+^/CD3^+^/B220^−^/CD8^+^. Active proliferating and Active Central Memory (CM) CD8^+^ T cells were identified as Ki-67^+^/CD62L^+^, respectively negative or positive for CD44. Naïve and DN CD8^+^ T cells were identified as Ki-67^−^/CD69^−^, respectively positive or negative for CD62L. Finally, the last three subsets were commonly identified as Ki-67^−^/CD69^+^, and divided into Active Naïve CD8^+^ T cells CD62L^+^/CD44^−^, Active CM CD8^+^ T cells CD62L^+^/CD44^+^, and Active DN CD8^+^ T cells CD62L^−^ /CD44^+^. **(C)** Histogram showing DN, Active CM, Proliferating CM CD8^+^ T cell subsets at 0 h, 24 h, or 7 days p.v. in the pLN. (n=3 for each group). Data are presented as mean ± SD. One-way ANOVA followed by Bonferroni correction for multiple comparisons (* p < 0.05, ** p < 0.01, *** p < 0.001 **** p<0.0001).

